# Sex differences in seizure at presentation in glioma population

**DOI:** 10.1101/718791

**Authors:** Sara Ranjbar, Barrett J Anderies, Kyle W Singleton, Sandra K Johnston, Cassandra R Rickertsen, Akanshka Sharma, Alyx B Porter, Maciej M Mrugala, Leland S Hu, J Ross Mitchell, Joshua B Rubin, Kristin R Swanson

## Abstract

Seizures are common presenting symptoms of primary brain tumors. Mechanisms of epileptogenesis are still unknown and are believed to be multifactorial. Previous studies have indicated correlation of seizure with tumor location. Recent investigations of our group have shown image-based parameters have sex-specific implications for patient outcome. In this retrospective study, we examined the association of tumor location with the probability and risk of seizure in male and female glioma patients.

## I. Introduction

Seizure is a common symptom of brain tumors and has been reported in 30-60% of GBM patients, in which two-third of seizures occurred at presentation [1]. Most tumor-associated seizures are initially focal, although secondary generalization may occur quickly and mask the occurrence of the initial focal seizure [2]. Low tumor grade and cortical tumor location have been suggested as the main risk factors for epilepsy [2]. Recent investigations of our group [3, 4] have demonstrated that image-based parameters have sex-specific implications for patient outcome. In this work, we investigated the association of seizure at presentation with tumor presence in brain structures in male and female high-grade glioma patients.

## II. Methods

We selected adult patients with glioma diagnosis (any grade), known status of symptomatic seizure, and available pretreatment T2-FLAIR or T2, and post gadolinium T1 images (T1GD). Table 1 describes our cohort. We co-registered the images (Figure 1A) with Harvard-Oxford probabilistic cortical and subcortical atlases [5]. We lateralized the cortical atlas by bisecting the mask into left and right hemispheres. The subcortical atlas contained several large regions of interest (ROIs) for the cortical structures including left/right white matter, gray matter, cerebrospinal fluid, and brain stem. We excluded gray and white matter ROIs from the subcortical mask and combined the remaining ROIs with the cortical ROIs. We consulted two neurologists to define larger lobe-level ROIs. These regions included lateralized frontal, temporal, parietal, occipital, limbic lobes, deep brain structures and the lateral ventricles (Figure 1B). Next, we assessed the presence of T1GD enhancement and T2-FLAIR hyperintensity infiltration on atlas ROIs for all patients, divided our patient cohort into males and females, and calculated probability of seizure given infiltration (PR) and relative regional risk of seizure given infiltration (RRR) for each group as defined in (1):

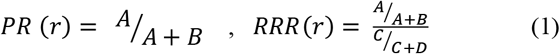

where r is a ROI and A, B, C, and D are the number of patients that satisfy the conditions in the cells of Table 2. We calculated the above separately for the abnormality seen on T1GD and FLAIR, and for males and females. Results were transformed into maps for visual comparison.

**Figure 1.**
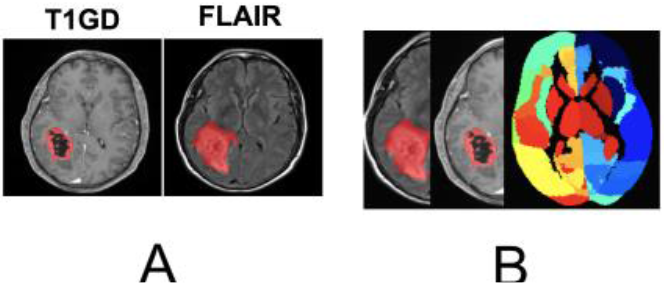
A. Example of T1GD and FLAIR images with the outline of abnormality on the images. B. images are co-registered with the atlas and the overlap of tumor abnormality on each image with atlas ROIs are calculated.

**TABLE 1.** PATIENT COHORT

|  | Seizure<br>Presenting<br>(SP) N = 60 | Non-Seizure<br>Presenting<br>(NSP) N=65 |  |
| --- | --- | --- | --- |
|  | Mean ± SD | Mean ± SD | P-value |
| Age | 52.4 ± 13.7 | 53.8 ± 14.3 | P = 0.583 |
| T1GD radius | 11.3 ± 6.25 | 19.3 ± 5.76 | P <0.001 |
| T2/FLAIR radius | 21.5 ± 7.44 | 27.5 ± 6.32 | P < 0.001 |
|  | N (%) | N (%) | P-value |
| Sex, Male | 45 (75%) | 34 (52%) | P = 0.009 |
| Grade 2 | 2 (100%) | 0 (0%) | - |
| Grade 3 | 9 (60%) | 6 (40%) | P = 0.35 |
| Grade 4 | 49 (45%) | 59(55%) |  |

**TABLE 2.**
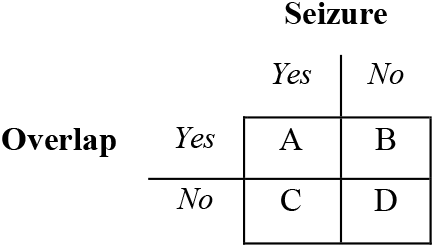
CONTINGENCY TABLE FOR RISK ASSESSMENT

## III. Results and discussion

We found higher incidence of seizure in males (58%) than in females (32%) regardless of the location of tumor. When we assessed the probability of seizure at each region without comparison with others, we found seizure to be slightly more probable for left hemisphere tumors in females (Figure 2, left panel). No such distinction was found in males (Figure 3, right panel). Relative regional risk maps shown in Figure 3 compare the probability of seizure in one region to that of all each region. In females, gliomas in the left frontal, parietal and temporal were riskier than others to present with seizure (Figure 3, left panel) which we think reveals that risk of seizure at presentation is specific to the location of glioma for females. In males (Figure 3, right panel), seizure appeared to be more agnostic to location, meaning regardless of where the lesion was located, all regions showed moderate risk of seizure (relative risk of 1, equivalent to 50% chance). Therefore, no specific region stood out more than others in having seizure as a presenting symptom.

**Figure 2.**
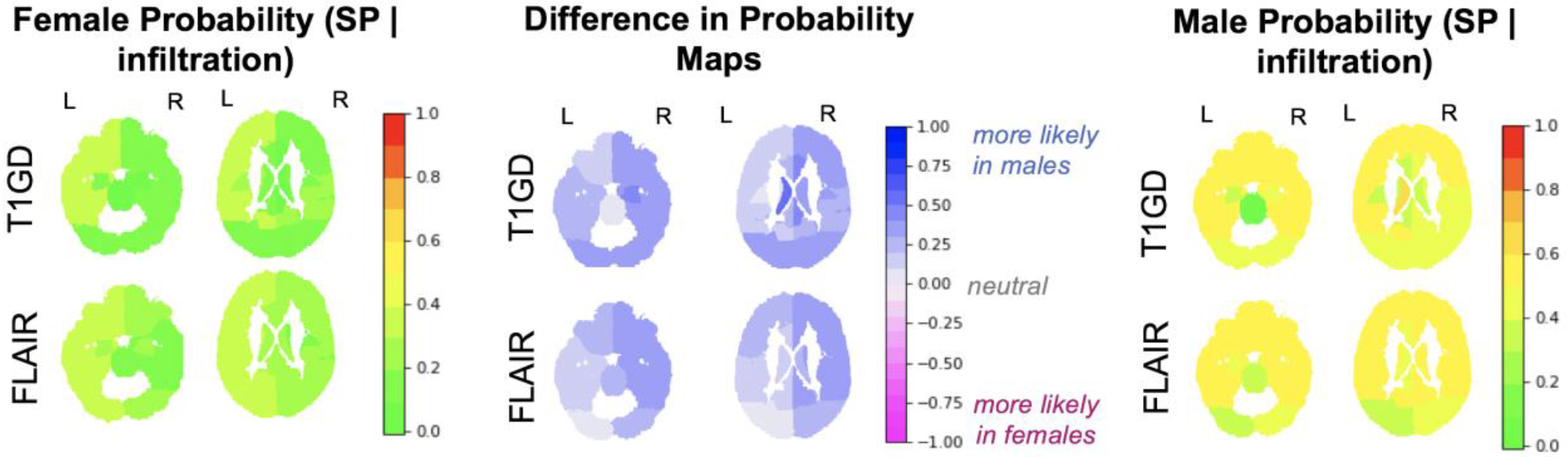
Probability of seizure in males (left) and females (right). Difference maps (middle) were generated by subtracting the female map from the male map.

**Figure 3.**
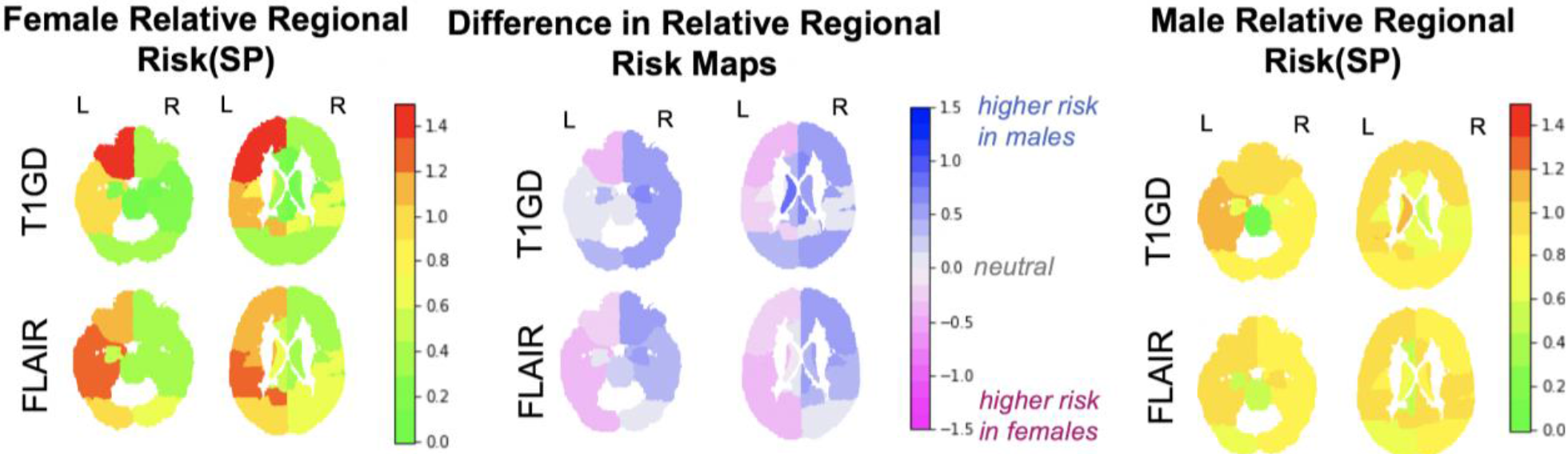
Relative regional risk of seizure in males (left) and females (right). Difference in the two maps (middle) were generated by subtracting the female map from the male map.

Our study has several limitations. Apart from small sample size, given the nature of a retrospective assessment we could not assure type of seizure in our patients. In the majority of these cases, seizure was self-reported without the possibility of a diagnosis of epilepsy. With these limitations in mind, our findings allow for the following conclusions. Overall, sensitivity to tumor infiltration was more pronounced for the regions in the left hemisphere in females. The specificity of risk in certain regions of female brains (red) may provide future guidance for seizure care management.

